# NODDI-DTI: extracting neurite orientation and dispersion parameters from a diffusion tensor

**DOI:** 10.1101/077099

**Authors:** Luke J. Edwards, Kerrin J. Pine, Nikolaus Weiskopf, Siawoosh Mohammadi

**Affiliations:** Department of Neurophysics, Max Planck Institute for Human Cognitive and Brain Sciences, Stephanstraβe 1a, 04103 Leipzig, Germany; Wellcome Trust Centre for Neuroimaging, UCL Institute of Neurology, UCL, 12 Queen Square, London, WC1N 3BG, UK.; Institut für Systemische Neurowissenschaften, Universitätsklinikum Hamburg-Eppendorf, Martinistraβe 52, 20246 Hamburg, UK

**Keywords:** diffusion MRI, NODDI, DTI, axonal density, axonal orientation dispersion

## Abstract

NODDI-DTI is a simplification of the NODDI model that, when its underlying assumptions are met, allows extraction of biophysical parameters from standard single-shell DTI data, permitting biophysical analysis of DTI datasets.

In contrast to the NODDI signal model, the NODDI-DTI signal model contains no CSF compartment, restricting its application to voxels with very low CSF partial-volume contamination. This simplification allowed derivation of simple analytical relations between parameters representing axon density, *ν*, and dispersion, *τ*, and DTI invariants (MD and FA) through moment expansion of the NODDI-DTI signal model. These relations formally allow extraction of biophysical parameters from DTI data. It should be emphasised that the NODDI-DTI model inherits the strong assumptions of the NODDI model, and so is nominally restricted to analysis of healthy brains.

NODDI-DTI parameter estimates were computed by applying the proposed analytical relations to DTI parameters extracted from the first shell of data, and compared to parameters extracted by fitting the NODDI-DTI model to (i) both shells (recommended) and (ii) the first shell (as for DTI) of data in the white matter of three different in vivo diffusion datasets. NODDI-DTI parameters estimated from DTI and NODDI-DTI parameters estimated by fitting the model to the first shell of data gave similar errors compared to two-shell NODDI-DTI estimates. The NODDI-DTI method gave unphysical parameter estimates in a small percentage of voxels, reflecting voxelwise DTI estimation error or NODDI-DTI model invalidity. In the course of evaluating the NODDI-DTI model it was found that diffusional kurtosis strongly biased DTI-based MD values, and so a novel heuristic correction requiring only DTI data was derived and used to account for this bias.

Our results demonstrate that NODDI-DTI is a promising model and technique to interpret restricted datasets acquired for DTI analysis with greater biophysical specificity, though its limitations must be borne in mind.

## Introduction

The white matter (WM) of the human brain consists of dense bundles of neuronal axons connecting its functional areas. Neural circuits thus formed allow these areas to work together as a coherent entity. Changes in WM impact these neural circuits, and are thus the subject of studies investigating pathology [1, 2, 3], and cognition and learning [4, 5].

Diffusion tensor imaging (DTI) [6] is, at present, the most commonly used method to observe WM changes in-vivo [4, 5, 7]. This is because DTI is simply implemented and time efficient while allowing robust extraction of complementary parameters (e.g. ‘fractional anisotropy’ (FA) and ‘mean diffusivity’ (MD) [8]) sensitive to microstructural WM changes [9], even in clinical contexts (see e.g. References [1, 3]). Despite its microstructural sensitivity, the model underlying DTI (gaussian anisotropic diffusion [6]) is unspecific to biological changes. Numerous studies show MD and FA change in white matter (e.g. due to learning a new skill [4] or the pathology of Alzheimer’s disease [2]), but cannot, in the absence of further information, distinguish e.g. changes in axon density from changes in axon arrangement.

In order to extract parameters of direct neurobiological relevance from diffusion MRI, we need biophysical models [10, 11, 9]. The majority of biophysical models (including the model introduced below) are ‘multicompartment models’. Such models assume voxelwise diffusion contrast arises from linear combination of diffusion signals from distinguishable water compartments. Numerous multicompartment models have been proposed (see e.g. References [12, 13, 14, 15, 16, 17, 18, 19, 20, 21]), but the complexity and lack of robustness of most of these models hinder their routine use in neuroscientific and clinical studies.

The NODDI (neurite orientation dispersion and density imaging) model [17] is a multicompartment model allowing robust and time-efficient extraction of maps of parameters representing neurite (in WM: axon) density and dispersion, and represents a trade-off between complexity, robustness, and acquisition-time duration. Robustness is achieved by fixing the values of several model parameters from earlier models [22, 18], reducing the number of fitted parameters. As a result, the amount of data required to invert the model is reduced, giving acquisition-time durations approaching those available in clinical settings [17]. NODDI is thus gaining popularity in diffusion application studies [23, 24, 15, 25, 26, 20, 27], though the potential of fixed parameters to lead to bias in the fitted parameters has been a source of criticism [15, 28, 21, 29].

Herein, we investigate a simplification of the NODDI model in which the CSF compartment (see below) is removed, and prove that there exist explicit relations between the remaining parameters of this model and MD and FA. We call this restricted version of the NODDI model ‘NODDI-DTI’ because it allows extraction of neurite orientation and dispersion parameters using MD and FA from DTI. The existence of these relations explains previously observed correlations between NODDI parameters and MD and FA [17, 30, 25, 31, 32]. Below, we investigate how accurately NODDI-DTI parameters extracted from DTI parameters in WM correspond to parameters fitted using the full NODDI-DTI model, and characterise limitations of the model and method.

## Theory

The NODDI signal model supposes three compartments: intraneurite water, extra-neurite water, and free water [17]. The biophysical parameters fitted in the model are neurite density (volume fraction of the intraneurite compartment), *ν*; a measure of neurite dispersion, *κ*; a vector giving the main neurite orientation; and a volume fraction accounting for partial-volume effects with free water (nominally CSF) [33, 34, 17]. An important fixed parameter is the intrinsic diffusivity of the intraneurite compartment, d = 1.7 × 10^−3^ mm^2^ s^−1^ [17]. The primary neurite orientation [17] is formally equivalent to the principal eigenvector of the diffusion tensor (DT) (see Appendix A), as previously observed empirically [35].

The model which we investigate here, which we call ‘NODDI-DTI’, is a reduced form of the NODDI model with no CSF volume fraction; this model has been previously observed to give reasonable estimates of *v* and *κ* from single-shell data [36]. For ease of computation in the following we use *τ* instead of *κ* as our measure of dispersion, where [15, 17, 18]

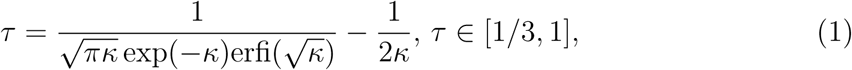

and erfi is the imaginary error function. *τ* ranges from 1/3 (isotropically distributed neurites) to 1 (perfectly aligned neurites)-increasing *τ* corresponds to increasing neurite alignment-and is the average of cos^2^(*ψ*) over the neurite distribution, where *ψ* is the angle between a given neurite and the main neurite orientation [15].

By expanding the NODDI-DTI signal model in moments, one can derive a corresponding DT [18]. As shown in Appendix A, appropriate combination of the eigenvalues of this DT allows expression of *ν* and *τ* in terms of MD and FA of this DT:

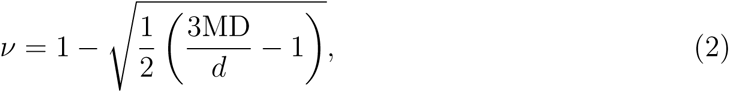

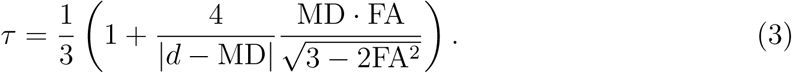

These relations are exemplified in Figure 1, and we note that Equation (2) has been previously derived independently [37].

**Figure 1:**
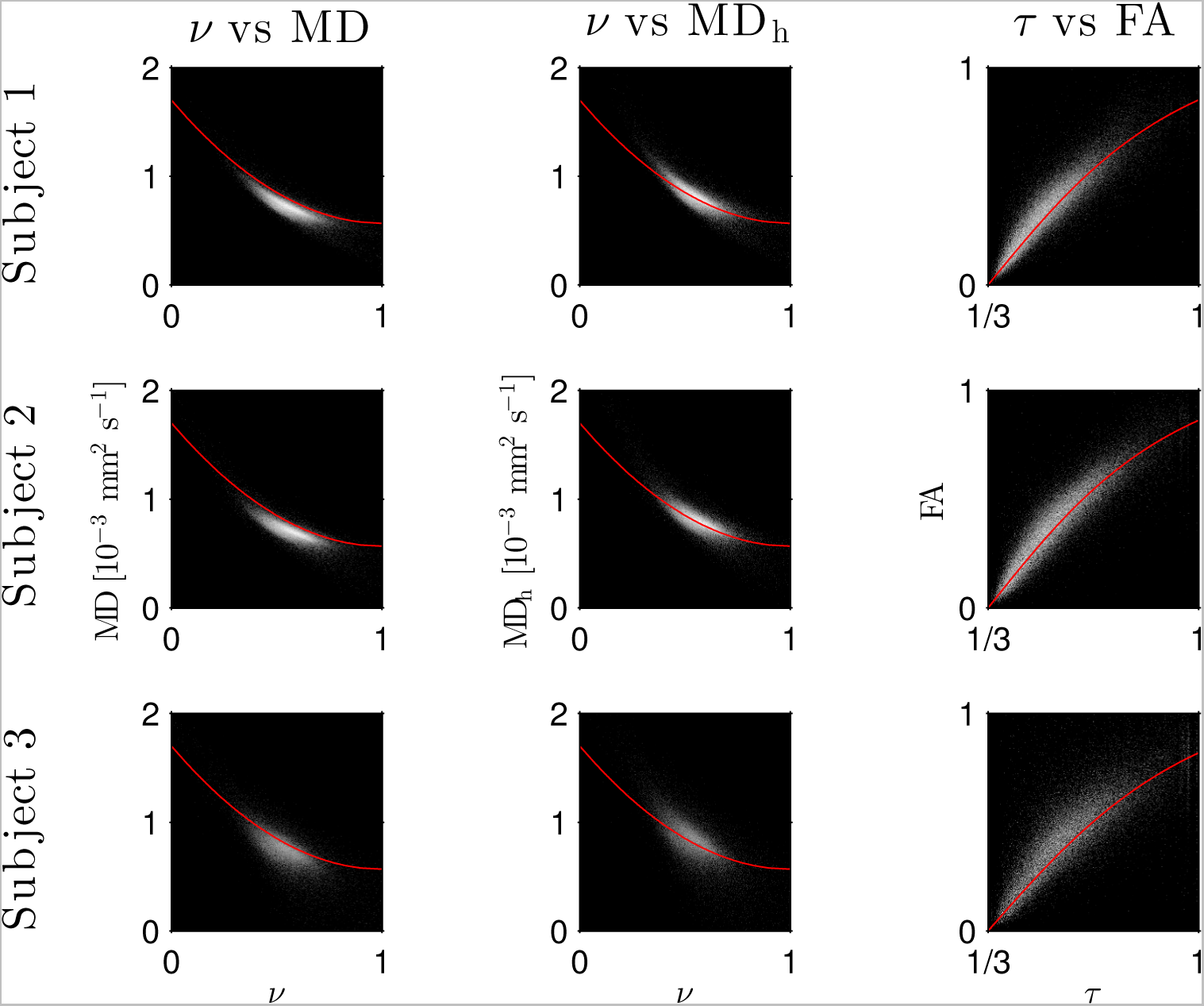
Log density scatter plots comparing DTI invariants (computed using only the low b-value shell of data) and NODDI-toolbox fitted parameters (fitted using both shells of data). Each row shows a different subject as labelled. Overlaid red lines in the first and second columns show values of *v* for given values of MD and MDh, respectively, computed using Eq. (2). The overlaid red lines in the third column show *τ* computed using Eq. (3) for given FA, with MD set to the mean value in the WM of each subject.

Eqs. (2) and (3) demonstrate a one-to-one mapping from (MD, FA) to (*v*, *τ*), demonstrating that, formally, NODDI-DTI parameters can be extracted from DTI data. We can predict domains within which MD and FA should lie if the NODDI-DTI model provides a valid representation: substituting *v* ∈ [0; 1] and *τ* ∈ [1/3; 1] into Eqs. (2) and (3) gives the domains

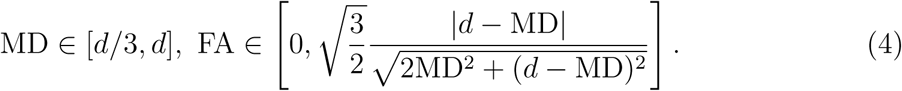

Exceptions will arise in practice either from DT measurement error or the invalidity of the NODDI-DTI model as a representation in a given voxel.

Estimation of a quantitative DT (i.e. the first moment of the diffusion signal) is non-trivial, as immediately apparent from examination of Figure 1: the main bulk of the MD values lies below the prediction of Eq. (2). A large part of this bias is due to neglecting higher order moments in estimating MD [38]. In order to reduce this bias, we define the heuristically corrected MD,

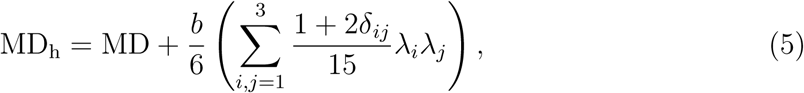

where λ_*i*_ is the *i*th eigenvalue of the measured DT and *δ*_*ij*_ is the Kronecker delta. Eq. (5), derived in Appendix B, pragmatically assumes that: only the first higher moment, diffusional kurtosis [39], contributes; the square of the apparent diffusion coefficient is uncorrelated with the apparent diffusional kurtosis; the mean diffusional kurtosis can be taken to be unity (approximately true over much healthy human brain WM [39, 40, 41]); and the effect of diffusional kurtosis on each individual eigenvalue is negligible. Figure 1 shows much closer agreement between the prediction of Eq. (2) and the experimental data when using MDh; further justification of this correction is given below.

Substituting Eq. (5) into Eq. (2) gives the relation used in the following to compute *ν* from DT invariants:

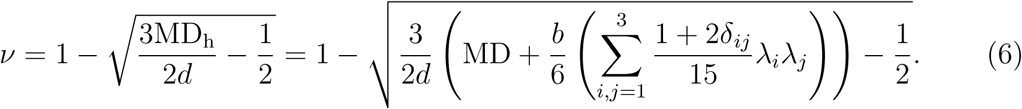

The effect of failing to correct for diffusional kurtosis is much less pronounced for FA [38], and preliminary experiments (data not shown) showed that applying diffusional kurtosis correction to only MD in Eq. (3) resulted in a modest increase in the number of unphysical *τ* parameter estimates. This latter observation can be explained using Eq. (4): whenever heuristic diffusional kurtosis correction leads to overestimation of MD, the upper bound for allowed FA values is artificially decreased, potentially leading to unphysical *τ* estimates. We therefore apply no correction to Eq. (3).

## Methods

### Data collection and preprocessing

All data were collected by scanning healthy volunteers in a MAGNETOM Tim Trio 3 T MRI system (Siemens AG, Healthcare Sector, Erlangen, Germany) as part of a study approved by a local ethics committee. In each case informed written consent was obtained prior to scanning.

The first two datasets (subject 1 and subject 2) used a 2D multiband spin-echo echo-planar imaging (EPI) sequence supplied by the Center for Magnetic Resonance Research, University of Minnesota [42, 43, 44]. Sequence parameters: field of view (FoV): 220 × 220 mm^2^; 81 slices, 1.7 mm isotropic resolution, echo time: TE = 112 ms, volume repetition time: TR = 4835 ms, partial Fourier factor: 6/8, multiband factor: 3, total 4 × 66 EPI images with 60 diffusion weighted images per shell using *b*-values of *b* = {1000; 2500} s mm^−2^, and 4 × 6 interleaved non-diffusion weighted (*b* = 0) images, 2 × phase encoding polarities (Anterior → Posterior / Posterior→Anterior).

The third dataset (subject 3) used a 2D spin-echo EPI sequence. Sequence parameters: FoV: 192×189 mm^2^, 63 slices, 2.0 mm isotropic resolution, TE = 100 ms, TR = 11700 ms, parallel imaging factor: 2, phase encoding polarity (Anterior → Posterior). Additional parameters for high *b*-value shell: partial Fourier factor: 6/8, total 111 EPI images with 100 diffusion weighted images with *b* = 2800 s mm^−2^ and 11 interleaved *b* = 0 images. Additional parameters for low *b*-value shell: no partial Fourier, total 110 EPI images with 100 diffusion weighted images with *b* = 800 s mm^−2^ and 10 interleaved *b* = 0 images.

In all cases (subjects 1–3) subject motion, eddy currents, and susceptibility distortions were corrected for using the ACID toolbox [45]; for details see References [46, 40, 47, 48]. Additionally for the multiband data (subjects 1 and 2), after the above corrections were applied, the corrected data from the two phase encoding directions were summed for use in subsequent analysis. This final step was unnecessary for subject 3.

### Parameter estimation and comparison

Parameters were estimated only in WM voxels determined to be largely unaffected by CSF or grey matter partial volume effects. This determination was made by thresholding at 50 % probability a WM probability map obtained by segmenting the first *b* = 0 image of each respective dataset in SPM 12 [49].

The ACID toolbox was used to compute FA, MD, and the eigenvalues of the DT from the low *b*-value shell of each dataset, and home-written SPM scripts were then used to generate *ν* and *τ* using Eqs. (6) and (3), respectively, from these DT parameters; we refer to these as the ‘NODDI-DTI method’ results. For subjects 1 and 2, the ACID toolbox was also used to simultaneously estimate the diffusion and kurtosis tensors [40], giving silver standard mean diffusivity estimates (MDDKI) less biased by the effects of diffusional kurtosis [38] allowing evaluation of the validity of Eq. 5. MDDKI was not computed for subject 3 as the second *b*-value was deemed too large to allow accurate estimation of the kurtosis tensor [39].

NODDI-DTI silver standard results were obtained by fitting both shells of data using the NODDI toolbox [17, 50] with CSF volume fraction fixed at zero, followed by conversion of *κ* into *τ* using a home-written SPM script implementing Eq. (1). We refer to these results as the ‘two-shell NODDI-toolbox fitted’ results, and they represent a silver-standard because two shells of data are suffcient to make inversion of multi-compartment signal models well-posed [51].

In order to investigate the magnitude of the differences between NODDI-DTI method- and the NODDI-toolbox fitted-results, fits were also made using the NODDI toolbox of the subset of the diffusion data used for the DTI fitting. The designation ‘one-shell NODDI-toolbox fitted’ distinguishes these results from the NODDI-DTI method- and two-shell NODDI-toolbox fitted-results. The NODDI toolbox has been shown previously to give reasonable results fitting single-shell data when CSF compartment fraction is fixed at zero [36].

Parameter estimate comparisons were quantified using means and standard deviations of the differences, visualised using Bland{Altman (BA) plots [52]. Voxels where Equations (3) and (6) gave unphysical parameter estimates (*v* ∈ [0; 1]; *τ* ∈ [1/3; 1]) were excluded from analysis.

## Results

Before examining the results of NODDI-DTI parameter extraction, we examine the heuristic correction of MD for diffusional kurtosis (Eq. (5)) used to calculate *ν*. Figure 1 and the BA plots in Figure 2 show that the heuristically corrected MD, MD_h_, is less biased compared to the uncorrected MD, justifying use of the correction. Numerical values (mean ± one standard deviation) of the differences in Figure 2 are for subject 1: 0:117 ± 0:040 (MD), 0:025 ±0:046 (MD_h_); and for subject 2: 0:119 ±0:041 (MD), 0:029 ±0:041 (MD_h_).

**Figure 2:**
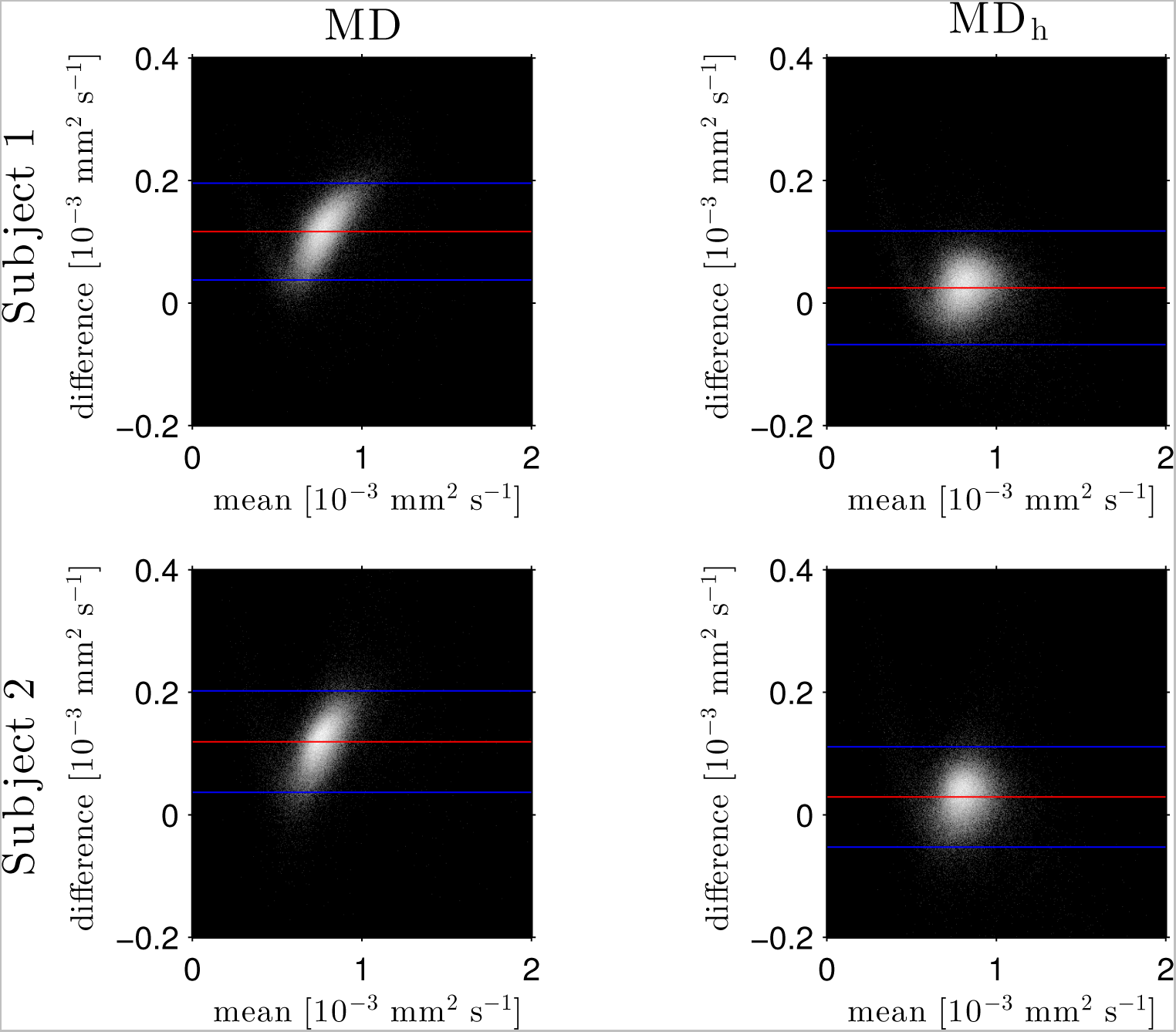
Log density Bland-Altman plots comparing MDDKI computed via simultaneous fit of the kurtosis tensor and DT using both shells of data, and mean diffusivity from a DT fit of the low-b-value shell without (MD, left) and with (MDh, right) heuristic diffusional kurtosis correction. Simultaneous fit of the kurtosis tensor and DT was performed as per Reference [40]. Differences are defined as MDDKI-(MD or MDh): Each row shows a different subject as labelled; results were not computed for subject 3 as the second b-value was deemed too large to allow accurate estimation of the kurtosis tensor [39]. Red lines show mean difference, blue lines show ± two standard deviations of the difference.

The similarity of parameters estimated using the NODDI-DTI method to the silver standard results can be seen in Figure 3, which shows parameter maps computed with each method, along with maps showing the differences between the parameter estimates. Differences between the parameter estimates are further presented in several complementary ways: BA plots in Figure 4 show general behaviour, plots of the means and standard deviations of the differences in Figure 5 compare this general behaviour across subjects, and the series of slices in Figures 6 and 7 show the behaviour of NODDI-DTI method parameter estimates throughout the brain. Subjects 1 and 2, measured using the same protocol, show the best agreement, but all three subjects behave similarly, demonstrating robustness of the NODDI-DTI method.

**Figure 3:**
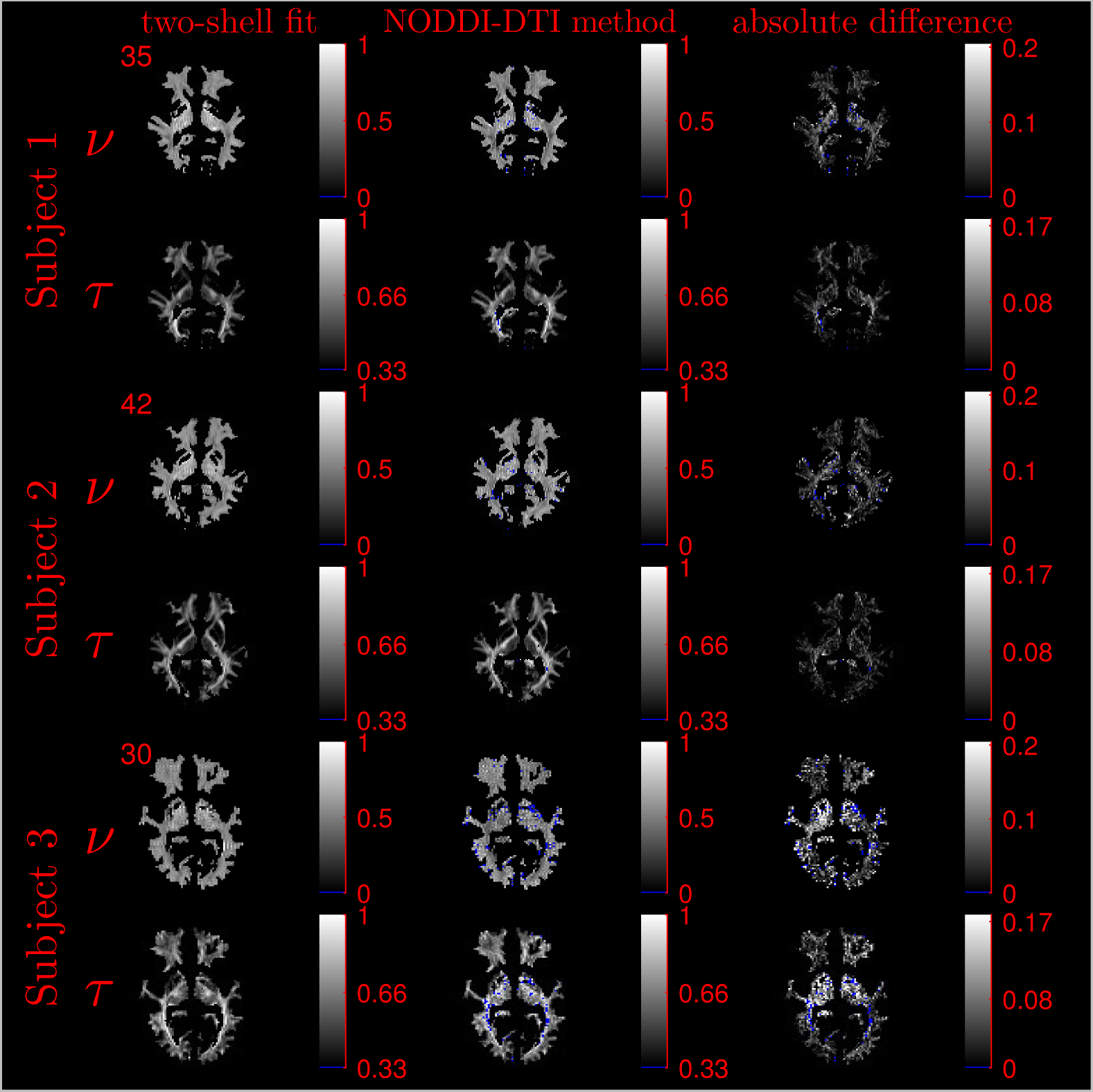
Comparison of maps of parameters computed using the NODDI-DTI method and fitting two-shells using the NODDI-toolbox. Voxels where the NODDI-DTI method gave an unphysical parameter estimate are shown in blue. Windows are as per the limits of the colour scales beside each map, and slice number is given at the top left of the row for each subject, to allow for cross-referencing with Figures 6 and 7.

**Figure 4:**
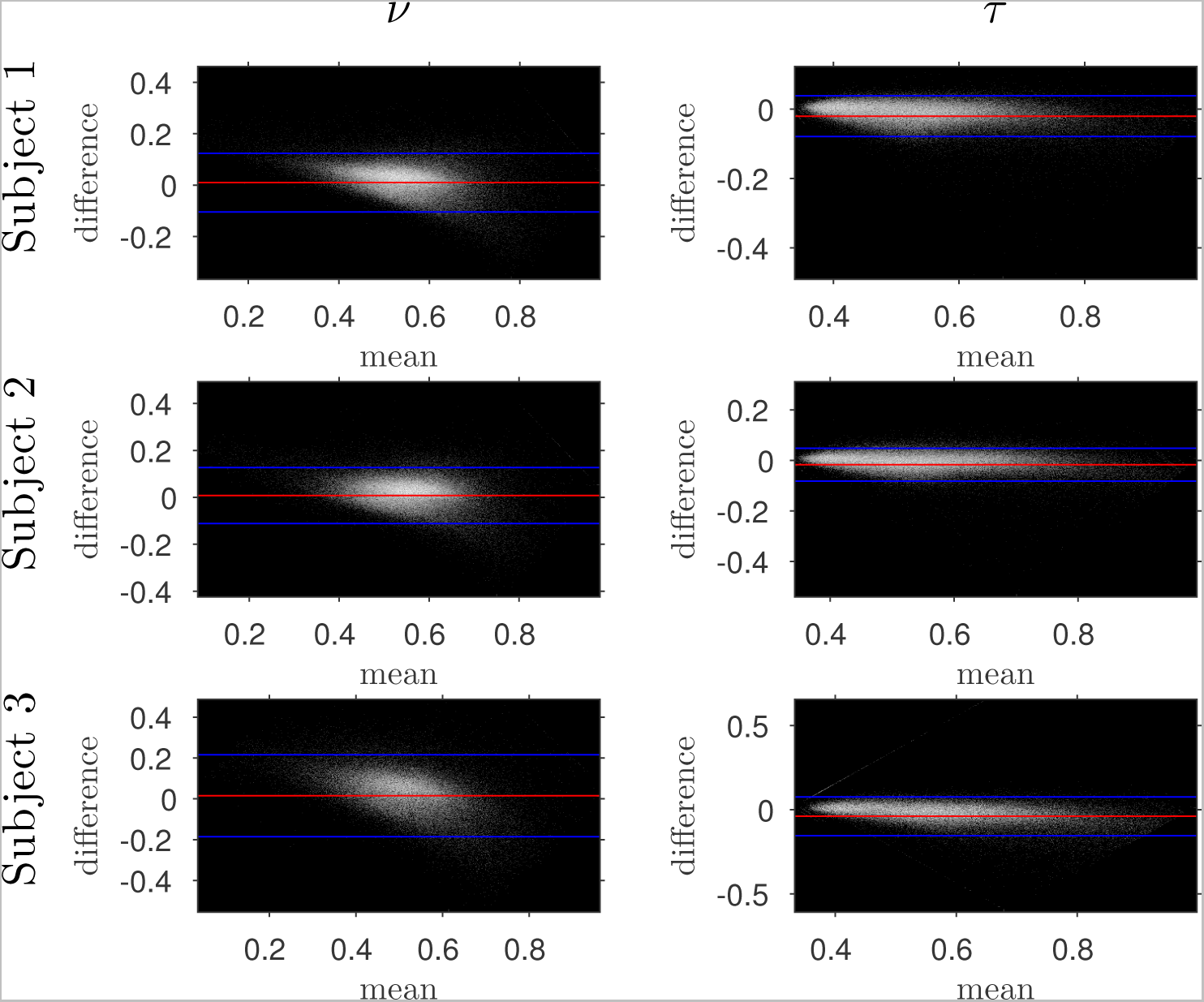
Bland-Altman plots comparing NODDI-DTI-method and two-shell NODDI-toolbox fitted results. Plotted is the log-density, and differences are defined as (two-shell NODDI-toolbox fitted parameter)-(NODDI-DTI method parameter): Red lines show mean difference, blue lines show ± two standard deviations of the difference; the numerical values of the means and standard deviations of the differences are given in Figure 5. Axis ranges show bounds of means and differences in each case.

**Figure 5:**
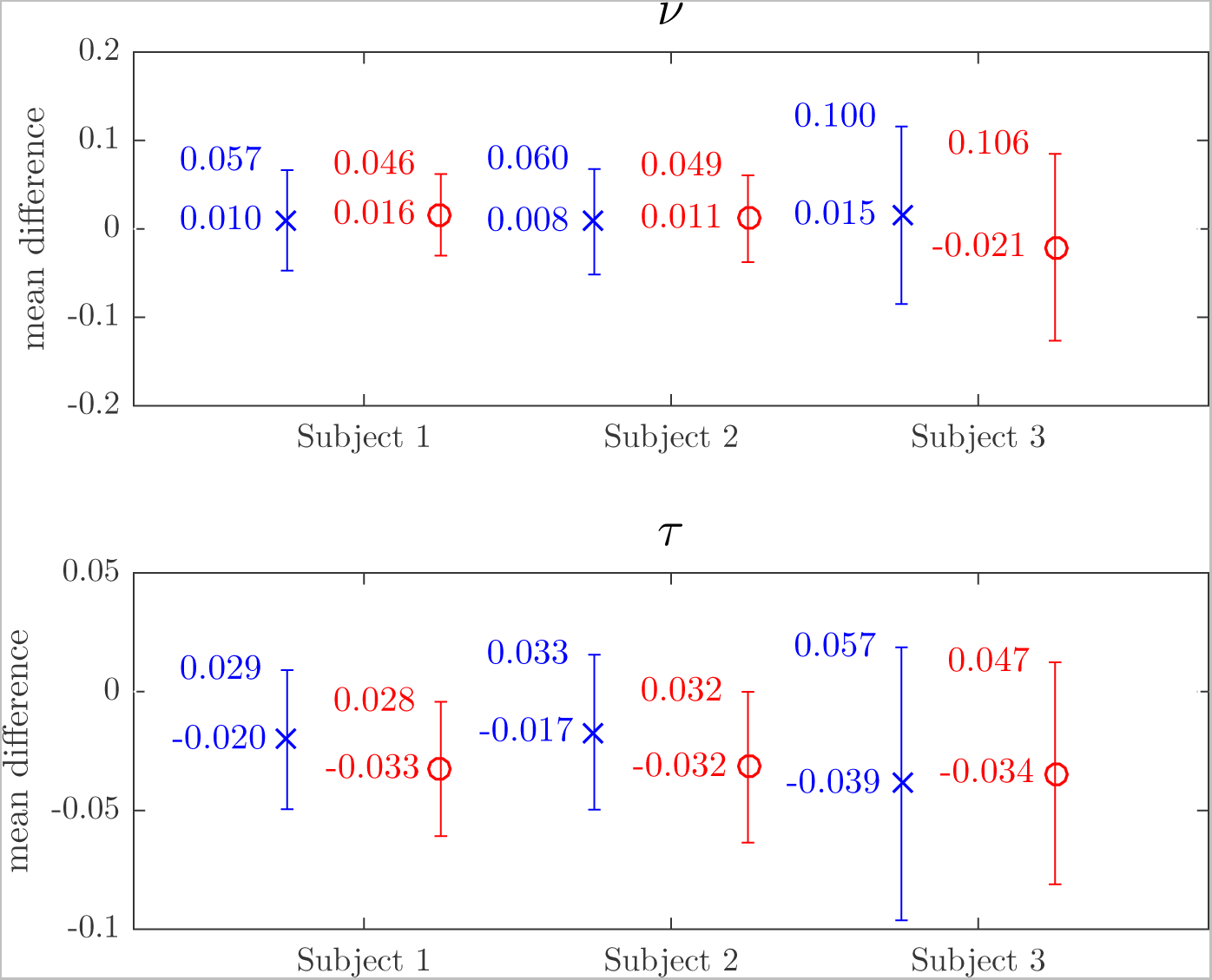
Plots of the mean differences between NODDI-DTI method and two-shell NODDI-toolbox fitted (×), and between one-shell NODDI-toolbox fitted (using the NODDI toolbox to fit the low b-value data) and two-shell NODDI-toolbox fitted (o) parameter estimates. Error bars show ± one standard deviation of the differences. Differences are defined as (two-shell NODDI-toolbox fitted parameter)-(estimated parameter): Numerical mean values are given beside each plotted point, and numerical values for the standard deviations are given beside the upper error bar.

**Figure 6:**
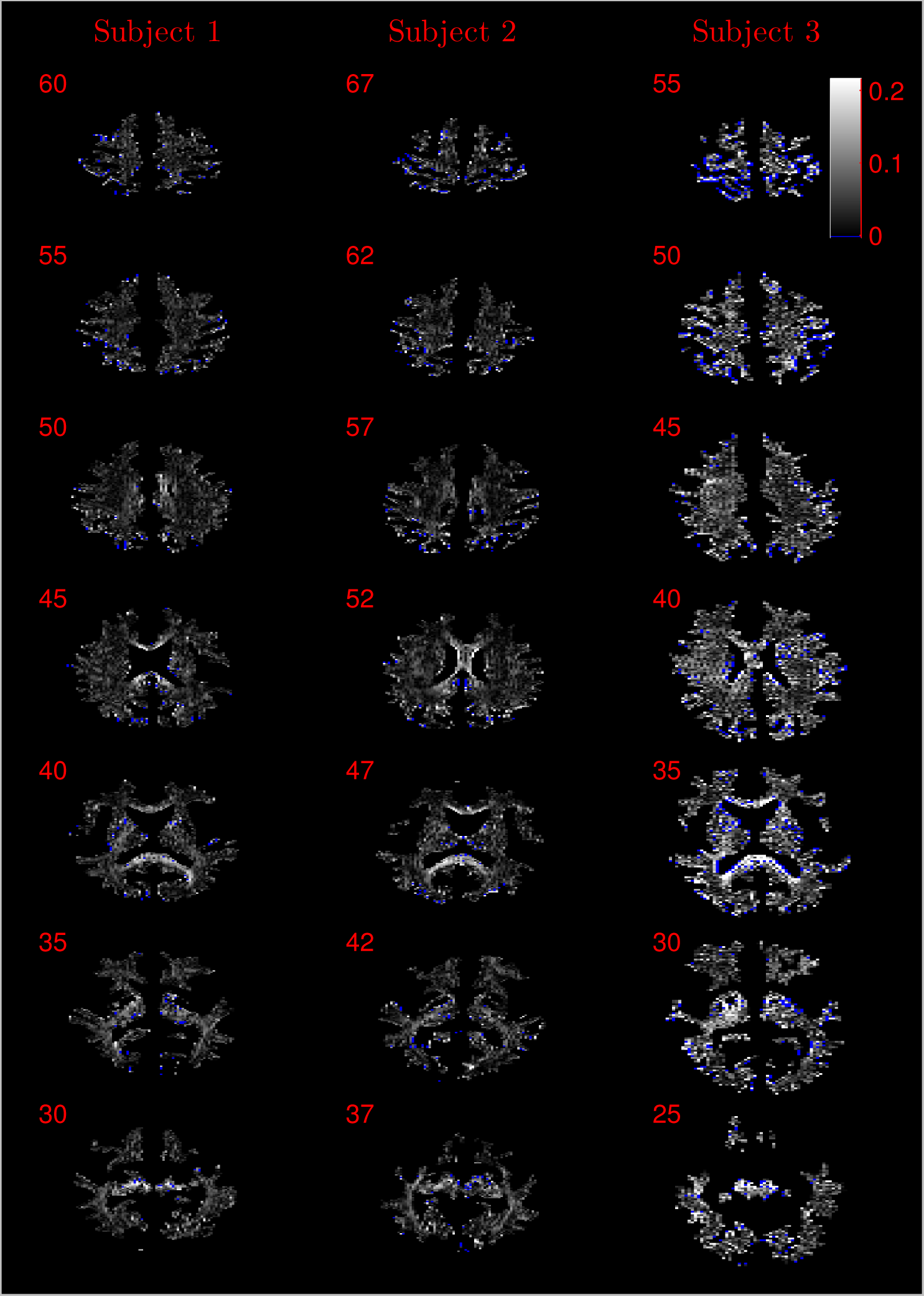
Absolute value maps of the difference between *v* computed using the NODDI-DTI method and fitting two-shells with the NODDI toolbox. Data from all three subjects are shown, and slice numbers are given for each row (slice) and column (subject). The extent of the colour scale at the top right shows the windowing for all slices. Blue denotes voxels where the NODDI-DTI method gave an unphysical parameter estimate.

**Figure 7:**
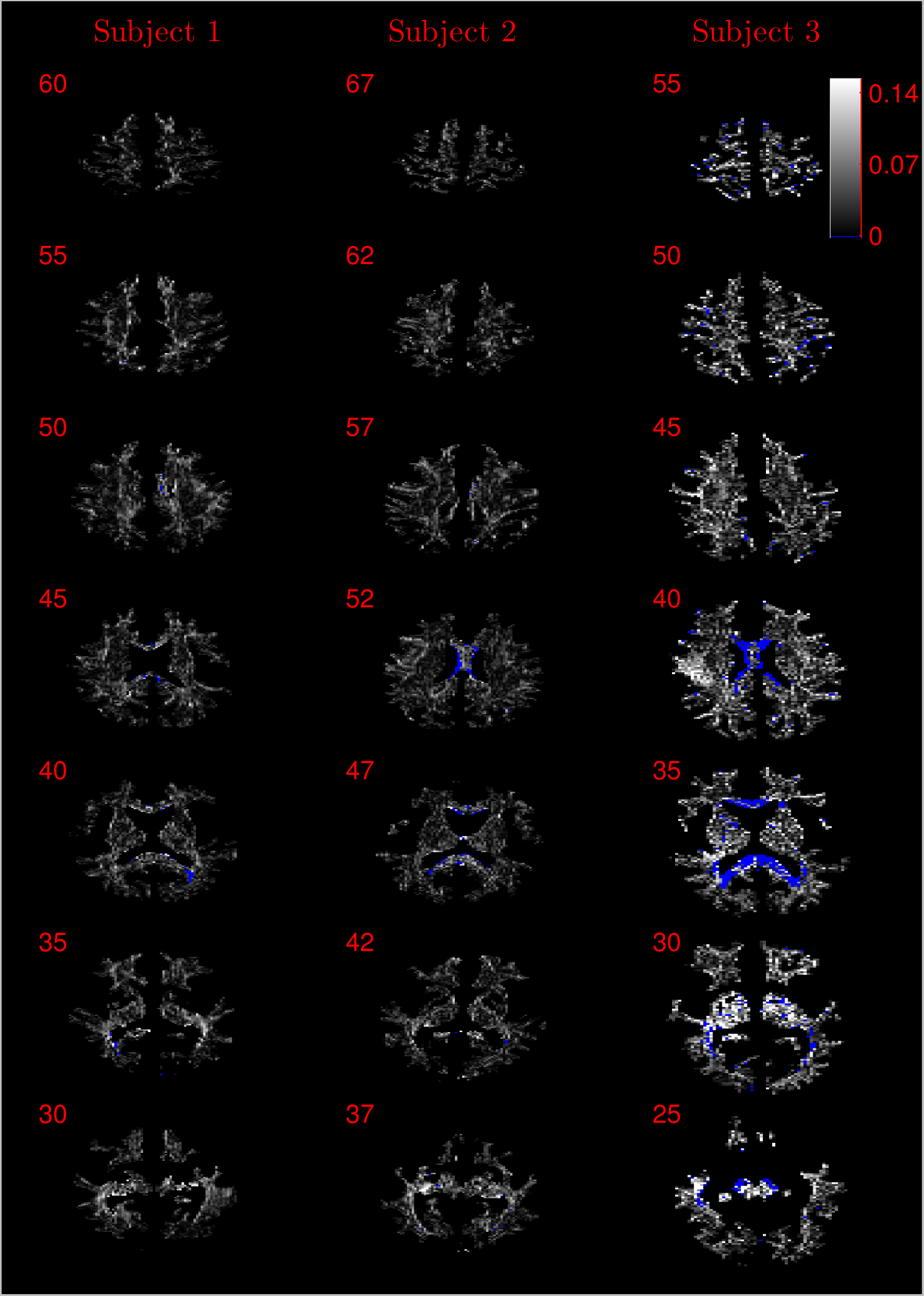
Absolute value maps of the difference between *τ* computed using the NODDI-DTI method and fitting two-shells with the NODDI toolbox. Data from all three subjects are shown, and slice numbers are given for each row (slice) and column (subject). The extent of the colour scale at the top right shows the windowing for all slices. Blue denotes voxels where the NODDI-DTI method gave an unphysical parameter estimate. The large number of unphysical *τ* values in the corpus callosum of subject 3 is discussed in the text.

Equations (3) and (6) gave unphysical parameter estimates for some voxels: for *ν* such voxels constituted 2.07 %, 2.19 %, and 6.83 % of the total WM voxels for subjects 1-3 respectively; for *τ* such voxels constituted 0.27 %, 0.74 %, and 5.39 %. The proportion of unphysical parameter estimates was higher for *ν* than *τ*, and greater for subject 3 than for subjects 1 and 2.

The magnitudes of the means and standard deviations of the differences between one- and two-shell NODDI-toolbox fitting are shown in Figure 5. One-shell NODDI-toolbox fitting gave stable fits in this case because the CSF compartment fraction was fixed at zero [36]. The NODDI-DTI method showed smaller mean differences and one-shell NODDI-toolbox fitting showed smaller standard deviations of the differences. Overall, however, both methods of parameter estimation were comparable, further demonstrating the validity of the NODDI-DTI method.

## Discussion

This work has demonstrated that, with caveats to be discussed below, parameters of potential neurobiological relevance can be extracted from DTI parameters using the NODDI-DTI relations, Equations (3) and (6). The improved interpretability thus gained would not only be applicable to future DTI studies, but also to existing DTI studies.

As an example of using NODDI-DTI to reinterpret existing DTI studies, we apply the method superficially to the study of Scholz, et al. [4], which demonstrated a statistically significant FA increase in WM “underlying the intraparietal sulcus” after participants learned to juggle [4]. Assuming no concomitant change in MD (as suggested by the authors not reporting any significant change), this FA increase could be interpreted, using Eq. (3), as an increase in *τ*, i.e. an increase in alignment of neuronal axons in this area with training. This result is much more specific than a change in FA, though we note that quantitative analysis would require reanalysis of the original data. A follow up study recording multi-shell data, combined with proper mechanistic analysis of the WM plasticity mechanisms, would allow investigation of this effect in more detail.

Unfortunately, the NODDI-DTI method did not always give physically plausible parameter estimates for the datasets studied herein. We posit four overlapping explanations for these unphysical estimates, related to NODDI-DTI assumptions.

Assumption 1: DTI parameters can be accurately estimated from the diffusion signal. Errors in DTI parameter estimation will lead to errors in parameters estimated using Equations (3) and (6), potentially giving unphysical parameter estimates. The *b*-values used likely resulted in overestimates of FA in regions of high anisotropy due to poor estimates of the low DT eigenvalues [8, 53], explaining why many of the unphysical *τ* estimates were found in the highly anisotropic corpus callosum (Figure 7). This hypothesis is bolstered by the results of subject 3: a lower *b*-value increased the proportion of unphysical estimates, especially in highly anisotropic regions.

Assumption 2: CSF can be ignored in voxels with a high probability of being WM. NODDI-DTI could give unphysical parameter estimates whenever a voxel contains a significant amount of CSF: the high difiusivity of CSF [17] can take MDh outside the limits of NODDI-DTI (Eqs. (4)). Figures 6 and 7 show that many of the voxels where NODDI-DTI gave unphysical parameter estimates are close to the edge of the WM mask, in line with partial volume effects being important. The larger voxel size used for scanning subject 3 means that partial volume effects are more prominent, partially explaining the poorer *ν* performance in this case. Because CSF volume fraction was fixed at zero in our NODDI-toolbox fits, residual partial volume effects may also have affected these parameter estimates.

Assumption 3: MD can be heuristically corrected for diffusional kurtosis bias. Figure 1 shows that diffusional kurtosis affects our estimates of MD; such effects could take MD estimates out of the range of applicability for NODDI-DTI. While our heuristic correction (Eq. (5)) substantially mitigates this issue, it does not eliminate it. This is evident in the *ν* Bland-Altman plots (Figure 4), where the mean and standard deviation of the differences visibly vary with the mean *ν* estimate, implying (via Eq. (6)) residual correlation between the errors and the corrected MD.

Residual bias in the region around the corpus callosum of the *ν* difference maps in Figure 6 can be explained by incomplete diffusional kurtosis correction. Here the mean diffusional kurtosis is greater than unity [39], and the low DT eigenvalues are poorly estimated (see Assumption 1), meaning that the heuristic correction is not entirely valid.

Assumption 4: the NODDI-DTI model is a valid representation of the diffusion signal. Assumptions underlying the NODDI-DTI signal model can be considered overly restrictive: the model cannot formally represent WM voxels containing perpendicularly crossing fibre bundles [54], and it is a subject of controversy as to whether intra-and extraneurite intrinsic difiusivities can be taken to be equal [28]. With respect to the latter point, NODDI-DTI will be unrepresentative whenever a voxel contains a significant amount of iron (e.g. through partial voluming with iron rich grey matter nuclei): susceptibility effects lower diffusivities artificially [55], and so lower *d*, assumed to be a fixed value in NODDI-DTI. In voxels where the NODDI-DTI signal model is unrepresentative, the NODDI-DTI method may give unphysical parameter estimates. Such failures will not be immediately apparent in the NODDI-toolbox fitted parameters because constraints in the fitting procedure mean parameters outside the physical range can never be returned, regardless of the model’s biological implausibility in a given voxel.

The greater number of unphysical NODDI-DTI method parameter estimates for *v* as compared to *τ* can be explained by *ν* estimation being more sensitive to partial volume and diffusional kurtosis effects. This is borne out by the locations of the failures (Figure 6): mainly either close to the edge of the WM mask (implying partial volume effects), or in regions of high anisotropy (implying residual diffusional kurtosis effects).

Pathology could further undermine the assumptions underlying NODDI-DTI: pathological processes can lead to free water located far from CSF compartments [56], can affect mean kurtosis values [57], and could affect the ‘true’ value of *d* [15].

NODDI-DTI could be improved and made more appropriate for clinical studies through investigation of the following points. Unphysical parameter estimates could be pragmatically eliminated by constraining DT fitting using Eqs. (4) (appropriately corrected using Eq. (5)). Estimates of CSF volume fraction could be incorporated into NODDI-DTI without requiring extra data acquisition using the free water elimination method [56, 34, 58]. Known values of mean diffusional kurtosis in the brain [39, 40, 41] could be used to construct mean diffusional kurtosis Bayesian priors [59, 51], or diffusional kurtosis corrected MD and FA could be measured directly using time-effcient methods [60, 61, 62].

We finish by providing practical recommendations for a minimal NODDI-DTI acquisition scheme. Results at *b* = 1000 mm^2^ s^−1^ were reasonable, and we would recommend this as a lower *b*-value bound; *b* = 800 mm^2^ s^−1^ unfortunately gave many unphysical parameter estimates in the corpus callosum. An upper bound on *b*-value comes from ensuring diffusional kurtosis does not constitute the majority of the diffusion contrast. Eq. (B.3) shows that (assuming the apparent diffusional kurtosis is approximately unity [39, 40, 41]) choosing *b* ≪ 6/*d* ≈ 3500 mm^2^s^−1^ means that the DT dominates diffusion contrast, giving an upper b-value bound. High resolution acquisitions which maintain good signal to noise ratio [53] are recommended to reduce partial volume effects. Accurate DT estimation requires measurement of at least 30 distinct diffusion directions [6]; we recommend at least this number for application of NODDI-DTI, although the lowest number of orientations tested was 60.

## Conclusions

We have estimated biophysical parameters representing neurite density and dispersion with reasonable accuracy from diffusion tensor parameters extracted from single-shell diffusion data. Heuristic kurtosis correction of MD was necessary to remove diffusional kurtosis bias; use of corrections such as that derived here could improve other analyses of single-shell diffusion data requiring quantitative MD estimates.

NODDI-DTI has the potential to open up two new opportunities: (a) more specific neurobiological interpretation of observed microstructural changes in DTI data (including interpretation of existing datasets), and (b) simple and time effcient estimation of biophysical parameters from smaller diffusion datasets, despite limitations due to the underlying model and difficulties estimating accurate diffusion tensors.

## Acknowledgements

The research leading to these results has received funding from the European Research Council under the European Union’s Seventh Framework Programme (FP7/2007-2013)/ ERC grant agreement no 616905. This project has received funding from the European Union’s Horizon 2020 research and innovation programme under the Marie Sk lodowska-Curie grant agreement No 658589. The Wellcome Trust Centre for Neuroimaging is supported by core funding from the Wellcome Trust 0915/Z/10/Z.

## Appendix A. Derivation of NODDI-DTI relations (Eqs. (2) and (3))

### Appendix A.1. Diffusion tensor of the NODDI-DTI signal model

We begin derivation of the NODDI-DTI relations by deriving the DT arising from the NODDI-DTI signal model. This derivation is similar to that of Reference [18], where the DT of a precursor to the NODDI-DTI model [22] was derived.

The normalised signal arising from the NODDI signal model can be written [17]

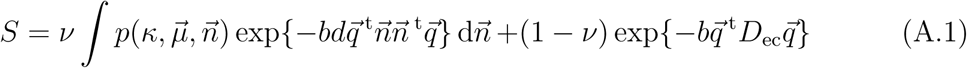

where the first term represents the intraneurite water compartment with diffusivity *d* parallel to the neurite and zero perpendicular to it; the second term represents the extraneurite water compartment; arrows denote normalised vectors;·^t^ denotes transposition; 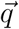 is the diffusion gradient vector; *ν* represents neurite density; and

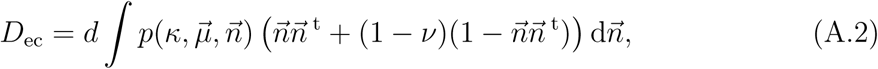

the DT of the extraneurite compartment. The form of the extraneurite DT arises from assuming that: the diffusivity of the extraneurite space in the absence of neurites is equal to the intraneurite diffusivity along the direction of the neurite [17], the neurites reduce the diffusivity in a long-time-limit tortuous manner [17], and extracellular water is in fast exchange among all neurite orientations [21]. The probability density

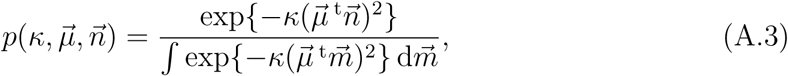

is a Watson distribution giving the distribution of neurites about main orientation 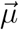 with dispersion parameter *κ* [17]. Isotropically distributed neurites correspond to *κ* = 0; neurites perfectly aligned along 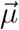 correspond to *κ → ∞*

Eq. (A.1) can be equated with an expansion of the normalised diffusion signal in *b* [63],

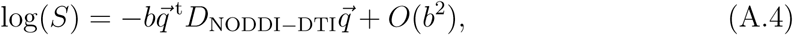

such that the DT can be extracted by inspection from

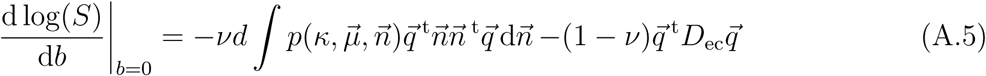

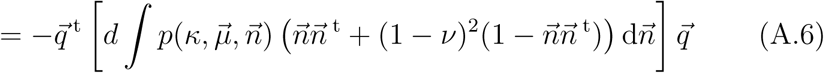

as

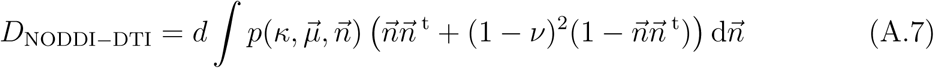

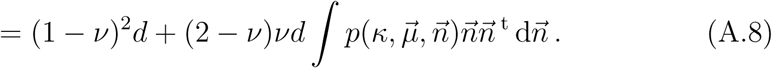

The integral appearing on the right-hand side of Eq. (A.8) is given by [18]

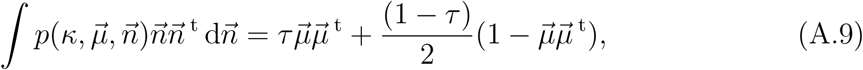

where *τ* is defined in Eq. (1). Inserting Eq. (A.9) into Eq. (A.8) gives

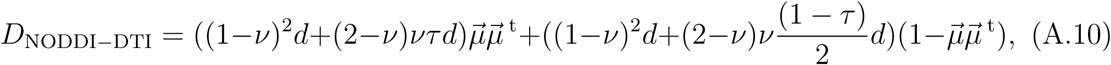

from which, by inspection, the largest eigenvalue (corresponding to an eigenvector co-linear with the main neurite orientation) is

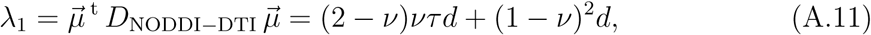

and the other two eigenvalues are degenerate (with respective eigenvectors arbitrarily defined in the plane perpendicular to 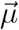):

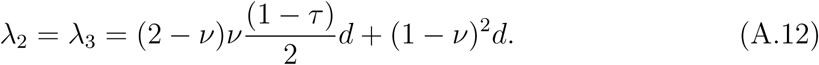

Because, λ_1_ ≥ λ_2_ when the primary eigenvector of *D*_NODDI-DTI_ is well-defined (i.e. when *D*_NODDI-DTI_ is not isotropic), this eigenvector is formally equivalent to the main neurite orientation, as observed empirically [35].

### Appendix A.2. Relation of *v* to MD (Eq. (2))

MD is defined in terms of the eigenvalues of a DT as [6]

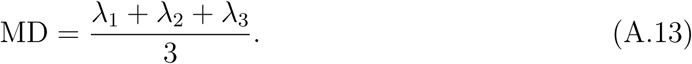

Inserting the eigenvalues from Eqs. (A.11) and (A.12) results in

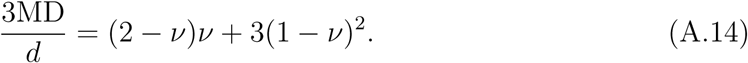

Solving this quadratic equation for *v* one obtains

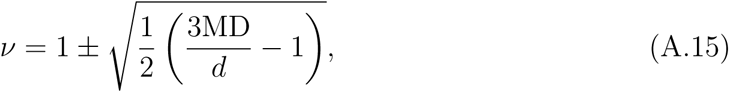

where the sign ambiguity is resolved by recalling *ν* ≤ 1 to give Eq. (2).

### Appendix A.3. Relation of *τ* to MD and FA (Eq. (3))

A convenient definition of FA in terms of the eigenvalues of a DT is [6]:

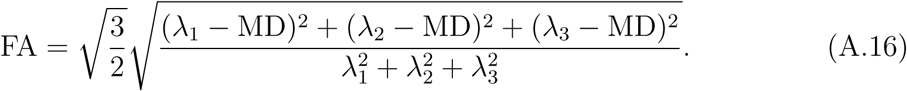

Because the eigenvalues are linear functions of *τ* (Eqs. (A.11) and (A.12)) and there is symmetry between them, it is convenient to simplify this equation by solving for λ_2_ before proceeding further. Utilising the identities λ_1_ = 3MD− λ_2_ − λ_3_ (Eq. (A.13)) and λ_2_ = λ_3_ Eq. (A.12)), Eq. (A.16) becomes:

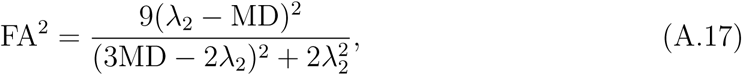

which can be rearranged into the quadratic equation

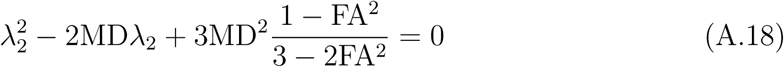

for which the solutions are:

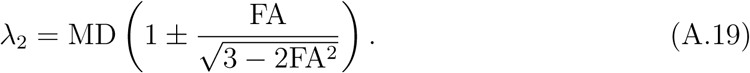

Eq. (A.19), reveals that all we must do to express *τ* in terms of MD and FA is (i) express *τ* in terms of MD and λ_2_ and then (ii) substitute Eq. (A.19) into the resulting expression.

Part (i) is achieved by substituting Eq. (2) into Eq. (A.12), then simplifying to give

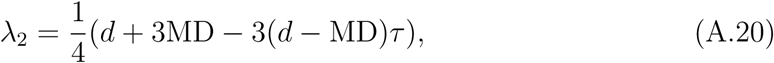

which, after rearranging for *τ*, reveals

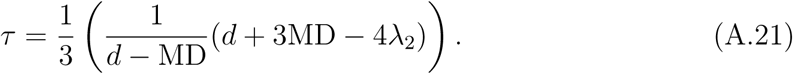

We can now perform part (ii): inserting Eq. (A.19) into Eq. (A.21) and simplifying gives the result

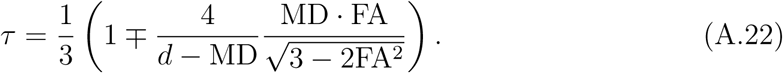

The sign ambiguity is resolved by recalling that *τ* ≥ 1/3 [15] and that both MD and FA are nonnegative, resulting in Eq. (3).

Eq. (3) is ill-defined at MD = *d* (the denominator of the second term goes to zero); we classify values at this point as ‘unphysical’ unless FA is also zero. This latter situation corresponds to the complete absence of fibres (as confirmed by inserting MD = *d* into Eq. (2)), and so *τ* is taken to equal its isotropic value, 1/3.

## Appendix B. Heuristic correction of MD for diffusional kurtosis (Eq. (5))

When diffusional kurtosis and higher order moments are zero, the normalised diffusion signal *S* is related to the apparent diffusivity, *D*_app_; by [64]

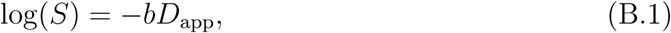

giving

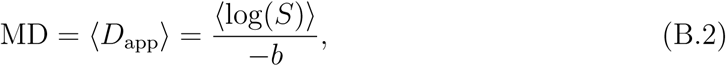

Where 〈·〉 denotes averaging over all diffusion directions.

The complicated microstructure of white matter requires higher order moments to represent the diffusion signal [63, 39, 38]. To the order of the diffusional kurtosis the normalised diffusion signal is

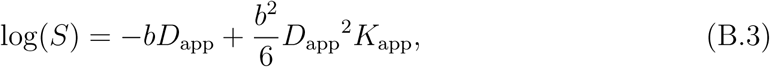

where *K*_app_ is the apparent diffusional kurtosis [63]. The effective mean diffusivity MD_eff_ derived from this signal (as per Eq. (B.2)) would be

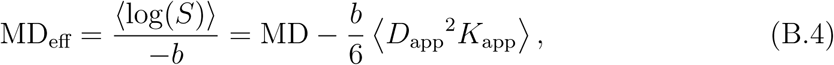

which differs from the true MD by a term which we call ‘diffusional kurtosis bias’. While good estimates of unbiased MD can be obtained from multi-*b*-value data [38], such extra data is not available for most DTI acquisitions, and so we derive and use an heuristic correction to mitigate diffusional kurtosis bias.

Defining the covariance of *D*_app_^2^ and *K*_app_:

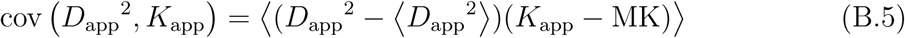

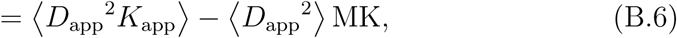

where MK = 〈*K*_app_〉; the mean kurtosis, we can write Eq. (B.4) in the form

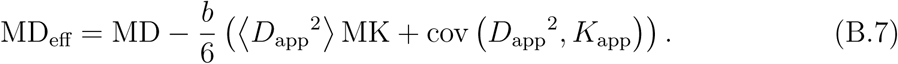

We pragmatically assert that cov (*D*_app_^2^; *K*_app_) = 0; i.e. assume that the apparent diffusivity and apparent diffusional kurtosis are uncorrelated. This assertion results in

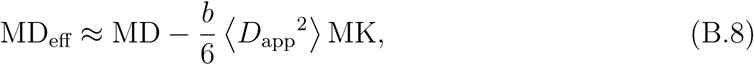

which is further simplified by assuming MK = 1 (true in much healthy WM [39, 40, 41]):

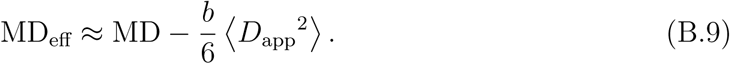

To compute the average in Eq. (B.9), we express *D*_app_ in components of the DT, *D*, and orientation vector, 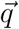, i.e. [39]

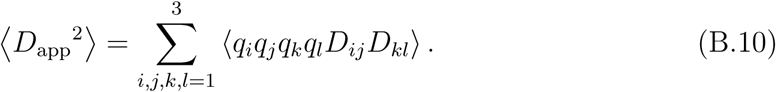

As we integrate over all 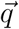 on the sphere, we can freely choose the basis of 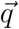. We thus choose the diagonal basis of *D*; simplifying Eq. (B.10) to:

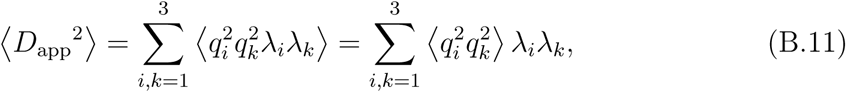

where λ*_i_* is the *i*th eigenvalue of *D* and is independent of orientation. Averages over the products of the components *q_i_*; evaluate to 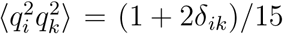, where *δ_ik_* is the Kronecker delta, thus

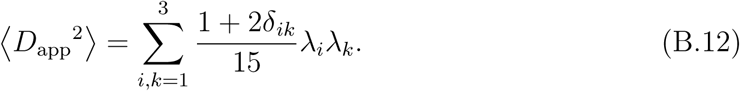

Inserting Eq. (B.12) into Eq. (B.9) and rearranging gives the heuristically corrected MD:

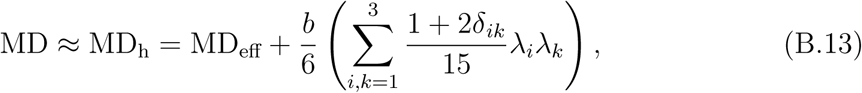

which becomes independent of diffusional kurtosis upon assuming the measured (diffusional kurtosis biased) eigenvalues can be substituted for the ‘true’ eigenvalues, giving Eq. (5).

## References

[1] M. Meinzer, S. Mohammadi, H. Kugel, H. Schiffbauer, A. Flöoel, J. Albers, K. Kramer, R. Menke, A. Baumgäartner, S. Knecht, C. Breitenstein, M. Deppe, Integrity of the hippocampus and surrounding white matter is correlated with language training success in aphasia, NeuroImage 53 (1) (2010) 283–290. doi: 10.1016/j.neuroimage.2010.06.004.

[2] J. Acosta-Cabronero, G. B. Williams, G. Pengas, P. J. Nestor, Absolute difiusivities define the landscape of white matter degeneration in Alzheimer’s disease, Brain 133 (2) (2010) 529–539. doi: 10.1093/brain/awp257.

[3] P. Freund, N. Weiskopf, J. Ashburner, K. Wolf, R. Sutter, D. R. Altmann, K. Friston, A. Thompson, A. Curt, MRI investigation of the sensorimotor cortex and the corticospinal tract after acute spinal cord injury: a prospective longitudinal study, The Lancet Neurology 12 (9) (2013) 873–881. doi: 10.1016/S1474-4422(13)70146-7.

[4] J. Scholz, M. C. Klein, T. E. Behrens, H. Johansen-Berg, Training induces changes in white-matter architecture, Nature Neuroscience 12 (11) (2009) 1370–1371. doi: 10.1038/nn.2412.

[5] R. J. Zatorre, R. D. Fields, H. Johansen-Berg, Plasticity in gray and white: neuroimaging changes in brain structure during learning, Nature Neuroscience 15 (4) (2012) 528–536. doi: 10.1038/nn.3045.

[6] D. K. Jones, Gaussian modeling of the diffusion signal, diffusion MRI, 2nd Edition, Academic Press, 2014, Ch. 5, pp. 87–104. doi: 10.1016/B978-0-12-396460-1.00005-6.

[7] R. Fields, Change in the brain’s white matter: The role of the brain’s white matter in active learning and memory may be underestimated, Science 330 (2010) 768–769. doi: 10.1126/science.1199139.

[8] C. Pierpaoli, P. Jezzard, P. Basser, A. Barnett, G. D. Chiro, Diffusion tensor MR imaging of the human brain, Radiology 201 (3) (1996) 637–648. doi: 10.1148/radiology.201.3.8939209.

[9] C. Beaulieu, The biological basis of diffusion anisotropy, diffusion MRI, 2nd Edition, Academic Press, San Diego, 2014, Ch. 8, pp. 155–183. doi: 10.1016/B978-0-12-396460-1.00008-1.

[10] K. K. Seunarine, D. C. Alexander, Multiple fibers: Beyond the diffusion tensor, Diffusion MRI, 2nd Edition, Academic Press, San Diego, 2014, Ch. 6, pp. 105–123. doi: 10.1016/B978-0-12-396460-1.00006-8.

[11] S. D. Santis, M. Drakesmith, S. Bells, Y. Assaf, D. K. Jones, Why diffusion tensor MRI does well only some of the time: Variance and covariance of white matter tissue microstructure attributes in the living human brain, NeuroImage 89 (2014) 35–44. doi: 10.1016/j.neuroimage.2013.12.003.

[12] G. J. Stanisz, G. A. Wright, R. M. Henkelman, A. Szafer, An analytical model of restricted diffusion in bovine optic nerve, Magnetic Resonance in Medicine 37 (1) (1997) 103–111. doi: 10.1002/mrm.1910370115.

[13] E. Fieremans, J. H. Jensen, J. A. Helpern, White matter characterization with diffusional kurtosis imaging, NeuroImage 58 (1) (2011) 177–188. doi: 10.1016/j.neuroimage.2011.06.006.

[14] E. Panagiotaki, T. Schneider, B. Siow, M. G. Hall, M. F. Lythgoe, D. C. Alexander, Compartment models of the diffusion MR signal in brain white matter: A taxonomy and comparison, NeuroImage 59 (3) (2012) 2241–2254. doi: 10.1016/j.neuroimage.2011.09.081.

[15] I. O. Jelescu, J. Veraart, V. Adisetiyo, S. S. Milla, D. S. Novikov, E. Fieremans, One diffusion acquisition and different white matter models: How does microstructure change in human early development based on WMTI and NODDI?, NeuroImage 107 (2015) 242–256. doi: 10.1016/j.neuroimage.2014.12.009.

[16] S. N. Jespersen, C. D. Kroenke, L. Østergaard, J. J. Ackerman, D. A. Yablonskiy, Modeling dendrite density from magnetic resonance diffusion measurements, NeuroImage 34 (4) (2007) 1473–1486. doi: 10.1016/j.neuroimage.2006.10.037.

[17] H. Zhang, T. Schneider, C. A. Wheeler-Kingshott, D. C. Alexander, NODDI: practical in vivo neurite orientation dispersion and density imaging of the human brain, NeuroImage 61 (4) (2012) 1000–1016. doi: 10.1016/j.neuroimage.2012.03.072.

[18] S. Jespersen, L. Leigland, A. Cornea, C. Kroenke, Determination of axonal and dendritic orientation distributions within the developing cerebral cortex by diffusion tensor imaging, IEEE Transactions on Medical Imaging 31 (1) (2012) 16–32. doi: 10.1109/TMI.2011.2162099.

[19] S. N. Sotiropoulos, T. E. Behrens, S. Jbabdi, Ball and rackets: Inferring fiber fanning from diffusion-weighted MRI, NeuroImage 60 (2) (2012) 1412–1425. doi: 10.1016/j.neuroimage.2012.01.056.

[20] M. Tariq, T. Schneider, D. C. Alexander, C. A. G. Wheeler-Kingshott, H. Zhang, Bingham-NODDI: Mapping anisotropic orientation dispersion of neurites using diffusion MRI, NeuroImage (2016) 207–223 doi: 10.1016/j.neuroimage.2016.01.046.

[21] E. Kaden, N. D. Kelm, R. P. Carson, M. D. Does, D. C. Alexander, Multi-compartment microscopic diffusion imaging, NeuroImage 139 (2016) 346–359. doi: 10.1016/j.neuroimage.2016.06.002.

[22] H. Zhang, P. L. Hubbard, G. J. Parker, D.C. Alexander, Axon diameter mapping in the presence of orientation dispersion with diffusion MRI, NeuroImage 56 (3) (2011) 1301–1315. doi: 10.1016/j.neuroimage.2011.01.084.

[23] J. P. Owen, Y. S. Chang, N. J. Pojman, P. Bukshpun, M. L. Wakahiro, E. J. Marco, J. I. Berman, J. E. Spiro, W. K. Chung, R. L. Buckner, T. P. Roberts, S. S. Nagarajan, E. H. Sherr, P. Mukherjee, the Simons VIP Consortium, Aberrant white matter microstructure in children with 16p11. 2 deletions, The Journal of Neuroscience 34 (18) (2014) 6214–6223. doi: 10.1523/JNEUROSCI.4495-13.2014.

[24] Y. S. Chang, J. P. Owen, N. J. Pojman, T. Thieu, P. Bukshpun, M. L. Wakahiro, J. I. Berman, T. P. Roberts, S. S. Nagarajan, E. H. Sherr, P. Mukherjee, White matter changes of neurite density and fiber orientation dispersion during human brain maturation, PLoS ONE 10 (6) (2015) e0123656. doi: 10.1371/journal.pone.0123656.

[25] F. Grussu, T. Schneider, H. Zhang, D. C. Alexander, C. A. Wheeler-Kingshott, Neurite orientation dispersion and density imaging of the healthy cervical spinal cord in vivo, NeuroImage 111 (2015) 590–601. doi: 10.1016/j.neuroimage.2015.01.045.

[26] Q. Wen, D. A. Kelley, S. Banerjee, J. M. Lupo, S. M. Chang, D. Xu, C. P. Hess, S. J. Nelson, Clinically feasible NODDI characterization of glioma using multiband EPI at 7 T, NeuroImage: Clinical 9 (2015) 291–299. doi: 10.1016/j.nicl.2015.08.017.

[27] J. S. W. Campbell, I. R. Leppert, S. Narayanan, T. Duval, J. Cohen-Adad, G. B. Pike, N. Stikov, Promise and pitfalls of g-ratio estimation with MRI, arXiv:1701. 02760 [physics, q-bio]. URL http://arxiv.org/abs/1701.02760.

[28] I. O. Jelescu, J. Veraart, E. Fieremans, D. S. Novikov, Degeneracy in model parameter estimation for multi-compartmental diffusion in neuronal tissue, NMR in Biomedicine 29 (1) (2016) 33–47. doi: 10.1002/nbm.3450.

[29] D. S. Novikov, J. Veraart, I. O. Jelescu, E. Fieremans, Mapping orientational and microstructural metrics of neuronal integrity with in vivo diffusion MRI, arXiv:1609. 09144 [physics, q-bio]. URL http://arxiv.org/abs/1609.09144.

[30] N. Kunz, H. Zhang, L. Vasung, K. R. O’Brien, Y. Assaf, F. Lazeyras, D. C. Alexander, P. S. Hüuppi, Assessing white matter microstructure of the newborn with multi-shell diffusion MRI and biophysical compartment models, NeuroImage 96 (2014) 288–299. doi: 10.1016/j.neuroimage.2014.03.057.

[31] A. R. Mayer, J. M. Ling, A. B. Dodd, T. B. Meier, F. M. Hanlon, S. D. Klimaj, A prospective microstructure imaging study in mixed-martial artists using geometric measures and diffusion tensor imaging: methods and findings, Brain Imaging and Behavior. (2016) 1–14. doi: 10.1007/s11682-016-9546-1.

[32] F. Deligianni, D. W. Carmichael, G. H. Zhang, C. A. Clark, J. D. Clayden, NODDI and Tensor-Based Microstructural Indices as Predictors of Functional Connectivity, PLOS ONE 11 (4) (2016) e0153404. doi: 10.1371/journal.pone.0153404.

[33] S. B. Vos, D. K. Jones, M. A. Viergever, A. Leemans, Partial volume effect as a hidden covariate in DTI analyses, NeuroImage 55 (4) (2011) 1566–1576. doi: 10.1016/j.neuroimage.2011.01.048.

[34] C. Metzler-Baddeley, M. J. O’Sullivan, S. Bells, O. Pasternak, D. K. Jones, How and how not to correct for CSF-contamination in diffusion MRI, NeuroImage 59 (2) (2012) 1394–1403. doi: 10.1016/j.neuroimage.2011.08.043.

[35] A. Daducci, E. J. Canales-Rodriguez, H. Zhang, T. B. Dyrby, D. C. Alexander, J.-P. Thiran, Accelerated microstructure imaging via convex optimization (AMICO) from diffusion MRI data, NeuroImage 105 (2015) 32–44. doi: 10.1016/j.neuroimage.2014.10.026.

[36] L. Magnollay, F. Grussu, C. A. Wheeler-Kingshott, V. Sethi, H. Zhang, D. Chard, D. H. Miller, O. Ciccarelli, An investigation of brain neurite density and dispersion in multiple sclerosis using single shell diffusion imaging, in: Proc. Intl. Soc. Mag. Reson. Med., 2014, abstract number 2048.

[37] B. Lampinen, F. Szczepankiewicz, J. Mårtensson, D. van Westen, P. C. Sundgren, M. Nilsson, Neurite density imaging versus imaging of microscopic anisotropy in diffusion MRI: A model comparison using spherical tensor encoding, NeuroImage 147 (2017) 517–531. doi: 10.1016/j.neuroimage.2016.11.053.

[38] J. Veraart, D. H. Poot, W. Van Hecke, I. Blockx, A. Vander Linden, M. Verhoye, J. Sijbers, More accurate estimation of diffusion tensor parameters using diffusion kurtosis imaging, Magnetic Resonance in Medicine 65 (1) (2011) 138–145. doi: 10.1002/mrm.22603.

[39] J. H. Jensen, J. A. Helpern, MRI quantification of non-gaussian water diffusion by kurtosis analysis, NMR in Biomedicine 23 (7) (2010) 698–710. doi: 10.1002/nbm.1518.

[40] S. Mohammadi, K. Tabelow, L. Ruthotto, T. Feiweier, J. Polzehl, N. Weiskopf, High-resolution diffusion kurtosis imaging at 3T enabled by advanced post-processing, Frontiers in Neuroscience 8 (427). doi: 10.3389/fnins.2014.00427.

[41] E. D. André, F. Grinberg, E. Farrher, I. I. Maximov, N. J. Shah, C. Meyer, M. Jaspar, V. Muto, C. Phillips, E. Balteau, Inuence of noise correction on intra-and inter-subject variability of quantitative metrics in diffusion kurtosis imaging, PLoS ONE 9 (4) (2014) 1–15. doi: 10.1371/journal.pone.0094531.

[42] S. Moeller, E. Yacoub, C. A. Olman, E. Auerbach, J. Strupp, N. Harel, K. Uğurbil, Multiband multislice GE-EPI at 7 Tesla, with 16-fold acceleration using partial parallel imaging with application to high spatial and temporal whole-brain fMRI, Magnetic Resonance in Medicine 63 (5) (2010) 1144–1153. doi: 10.1002/mrm.22361.

[43] K. Setsompop, B. A. Gagoski, J. R. Polimeni, T. Witzel, V. J. Wedeen, L. L. Wald, Blipped controlled aliasing in parallel imaging for simultaneous multislice echo planar imaging with reduced g-factor penalty, Magnetic Resonance in Medicine 67 (5) (2012) 1210–1224. doi: 10.1002/mrm.23097.

[44] J. Xu, S. Moeller, E. J. Auerbach, J. Strupp, S. M. Smith, D. A. Feinberg, E. Yacoub, K. Uğurbil, Evaluation of slice accelerations using multiband echo planar imaging at 3 T, NeuroImage 83 (2013) 991–1001. doi: 10.1016/j.neuroimage.2013.07.055.

[45] ACID toolbox for SPM.URL http://www.diffusiontools.com/.

[46] S. Mohammadi, H. E. Müoller, H. Kugel, D. K. Müuller, M. Deppe, Correcting eddy current and motion effects by affine whole-brain registrations: Evaluation of three-dimensional distortions and comparison with slicewise correction, Magnetic Resonance in Medicine 64 (4) (2010) 1047–1056. doi: 10.1002/mrm.22501.

[47] L. Ruthotto, H. Kugel, J. Olesch, B. Fischer, J. Modersitzki, M. Burger, C. H. Wolters, Diffeomorphic susceptibility artifact correction of diffusion-weighted magnetic resonance images, Physics in Medicine and Biology 57 (18) (2012) 5715. doi: 10.1088/0031-9155/57/18/5715.

[48] L. Ruthotto, S. Mohammadi, C. Heck, J. Modersitzki, N. Weiskopf, Hyperelastic Susceptibility Artifact Correction of DTI in SPM, Bildverarbeitung für die Medizin 2013, Informatik aktuell, Springer Berlin Heidelberg, 2013, pp. 344–349. doi: 10.1007/978-3-642-36480-8_60.

[49] SPM 12. URL http://www.fil.ion.ucl.ac.uk/spm/.

[50] NODDI toolbox v0.9. URL http://www.nitrc.org/projects/noddi_toolbox/.

[51] M. Taquet, B. Scherrer, N. Boumal, J. M. Peters, B. Macq, S. K. Warfield, Improved fidelity of brain microstructure mapping from single-shell diffusion MRI, Medical Image Analysis 26 (1) (2015) 268–286. doi: 10.1016/j.media.2015.10.004.

[52] J. M. Bland, D. G. Altman, Statistical methods for assessing agreement between two methods of clinical measurement, The Lancet 327 (8476) (1986) 307–310, originally published as Volume 1, Issue 8476. doi: 10.1016/S0140-6736(86)90837-8.

[53] D. K. Jones, P. J. Basser, \Squashing peanuts and smashing pumpkins": How noise distorts diffusion-weighted MR data, Magnetic Resonance in Medicine 52 (5) (2004) 979–993. doi: 10.1002/mrm.20283.

[54] B. Jeurissen, A. Leemans, J.-D. Tournier, D. K. Jones, J. Sijbers, Investigating the prevalence of complex fiber configurations in white matter tissue with diffusion magnetic resonance imaging, Human Brain Mapping 34 (11) (2013) 2747–2766. doi: 10.1002/hbm.22099.

[55] J. Zhong, R. P. Kennan, J. C. Gore, Effects of susceptibility variations on NMR measurements of diffusion, Journal of Magnetic Resonance 95 (2) (1991) 267–280. doi: 10.1016/0022-2364(91)90217H.

[56] O. Pasternak, N. Sochen, Y. Gur, N. Intrator, Y. Assaf, Free water elimination and mapping from diffusion MRI, Magnetic Resonance in Medicine 62 (3) (2009) 717–730. doi: 10.1002/mrm.22055.

[57] C. Guglielmetti, J. Veraart, E. Roelant, Z. Mai, J. Daans, J. Van Audekerke, M. Naeyaert, G. Vanhoutte, R. Delgado y Palacios, J. Praet, E. Fieremans, P. Ponsaerts, J. Sijbers, A. Van der Linden, M. Verhoye, Diffusion kurtosis imaging probes cortical alterations and white matter pathology following cuprizone induced demyelination and spontaneous remyelination, NeuroImage 125 (2016) 363–377. doi: 10.1016/j.neuroimage.2015.10.052.

[58] T. van Bruggen, H. Zhang, O. Pasternak, H.-P. Meinzer, B. Stieltjes, K. H. Fritzsche, Free-water elimination for assessing microstructural gray matter pathology - with application to Alzheimer’s disease, in: Proc. Intl. Soc. Mag. Reson. Med., 2013, abstract number 0790.

[59] D. C. Alexander, D. Zikic, V. Wottschel, J. Zhang, H. Zhang, A. Criminisi, Image quality transfer: exploiting bespoke high-quality data to enhance everyday acquisitions, in: Proc. Intl. Soc. Mag. Reson. Med., 2015, abstract number 0563.

[60] B. Hansen, T. E. Lund, R. Sangill, S. N. Jespersen, Experimentally and computationally fast method for estimation of a mean kurtosis, Magnetic Resonance in Medicine 69 (6) (2013) 1754–1760. doi: 10.1002/mrm.24743.

[61] B. Hansen, T. E. Lund, R. Sangill, S. N. Jespersen, Erratum: Hansen, Lund, Sangill, and Jespersen. Experimentally and computationally fast method for estimation of a mean kurtosis. Magnetic Resonance in Medicine 69:1754–1760 (2013), Magnetic Resonance in Medicine 71 (6) (2014) 2250–2250. doi: 10.1002/mrm.25090.

[62] B. Hansen, T. E. Lund, R. Sangill, E. Stubbe, J. Finsterbusch, S. N. Jespersen, Experimental considerations for fast kurtosis imaging, Magnetic Resonance in Medicine (2015) doi: 10.1002/mrm.26055.

[63] J. H. Jensen, J. A. Helpern, A. Ramani, H. Lu, K. Kaczynski, diffusional kurtosis imaging: The quantification of non-gaussian water diffusion by means of magnetic resonance imaging, Magnetic Resonance in Medicine 53 (6) (2005) 1432–1440. doi: 10.1002/mrm.20508.

[64] P. Basser, J. Mattiello, D. LeBihan, MR diffusion tensor spectroscopy and imaging, Biophysical Journal 66 (1) (1994) 259–267. doi: 10.1016/S0006-3495(94)80775-1.

